# Single Unit Analysis and Wide-Field Imaging Reveal Alterations in Excitatory and Inhibitory Neurons in Glioma

**DOI:** 10.1101/2021.02.23.432381

**Authors:** Brian J.A. Gill, Farhan A. Khan, Edward M. Merricks, Alexander R. Goldberg, Xiaoping Wu, Alexander A. Sosunov, Tejaswi D. Sudhakar, Jyun-you Liou, Athanassios Dovas, Guy M. McKhann, Peter Canoll, Catherine Schevon

## Abstract

Several studies have attributed the development of tumor-associated seizures to an excitatory-inhibitory imbalance, highlighting the importance of resolving the spatiotemporal interplay of different neuronal populations within the peritumoral microenvironment. We combined methods for microelectrode array recordings and single unit analysis during ictal events with wide-field optical mapping of pyramidal neurons in an *ex vivo* acute slice model of diffusely infiltrating glioma in Thy1-GCaMP6f mice. This approach allowed for characterization of excitatory and inhibitory populations across an extended peritumoral cortical region. As expected, measures of excitability were increased in tumor-bearing slices compared to control. This was accompanied by marked functional alterations in single units classified as fast-spiking interneurons, including reduced firing, altered timing with respect to excitatory firing, and deficits in surround inhibition. Inhibiting mTOR with AZD8055 reversed these glioma-induced changes to excitatory and inhibitory neuronal populations, suggesting a prominent role for functional mechanisms linked to mTOR activation.

## Introduction

Tumor-associated seizures (TAS) are the most common presenting symptom in patients with diffusely infiltrating gliomas^1^. Such events result from glioma-driven changes to neurons in the peritumoral microenvironment, including alterations in neuronal number^2–4^, neurotransmitter concentration^5, 6^, and ion channel expression^2, 7^. These changes disrupt the excitatory-inhibitory balance within the surrounding cortex, causing hyperexcitability and facilitating the development of epileptiform activity. Such aberrant neuronal firing can upregulate oncogenic pathways through both synaptic^8, 9^ and non-synaptic^10, 11^ mechanisms, highlighting the importance of studying this process.

We and others have shown that TAS arise from the peritumoral cortex surrounding gliomas^4, 12, 13^. This cortex contains glioma cells intermingled with non-neoplastic brain cells^14^, which include pyramidal neurons^8, 9, 15^ and parvalbumin-expressing (PV) inhibitory interneurons^3, 7, 15^. While previous studies have detailed how both excitatory^4, 6, 16, 17^ and inhibitory^3, 4^ neurons may be involved in TAS, few have examined the interplay between these two cell classes^2, 7^. A comprehensive understanding of how excitatory and inhibitory neuronal populations contribute to ictal events observed in tumoral epilepsy requires simultaneous observation and analysis of each population across an extended spatial region. This can be achieved with single unit analysis^18–20^ and cell type-specific genetically encoded calcium indicators^21^. Implementing this methodology in a model of TAS also allows for a greater understanding of how targeted drug therapy could affect these populations.

Herein, we combined our previously described methods for high density microelectrode array (MEA) recordings^12^ and single unit analysis during ictal events^18^, with simultaneous wide-field optical imaging of pyramidal neurons using an *ex vivo* acute slice model of diffusely infiltrating glioma in Thy1-GCaMP6f mice^13^. This approach allowed us to correlate local field potentials (LFP), multi-unit activity (MUA), and single unit activity with observations of excitatory activity captured through GCaMP imaging. We showed that increased excitability of pyramidal neurons in the peritumoral cortex coincided with dysfunctional inhibitory interneuron behavior. We pharmacologically perturbed this model by inhibiting mammalian target of rapamycin (mTOR), which has been implicated in both gliomagenesis^10, 11, 22^ and epileptogenesis^23–25^. Doing so enabled us to investigate the effects of mTOR inhibition on the crosstalk between excitatory and inhibitory neuronal populations over an extended spatial area of the peritumoral microenvironment.

## Results

### GCaMP transients coincide with bursts in LFP and MUA during epileptiform activity

Wide field optical imaging and dense MEA recordings were used to characterize the peritumoral regions of acute *ex vivo* slices obtained 25-30 days post-injection from a previously described murine model of diffusely infiltrating glioma, in which PDGFA+/p53-glioma cells were injected into Thy1-GCaMP6f mice. These mice expressed the calcium-sensitive green fluorescent protein in excitatory Thy1 positive pyramidal cells, typically found in layers 2, 3, and 5. Coronal slices obtained from these mice first underwent a 5-minute recording in artificial CSF (aCSF) followed by a 30-minute recording in zero-magnesium (zero-Mg^2+^) solution (Fig 1A) (Supplementary Movie 1). The recording was limited to this early time period as it best mimicked the glutamatergic environment observed in human glioma^6^. To determine whether GCaMP fluctuations correlated with electrophysiologic changes, we examined LFP line-length and MUA firing rate during GCaMP transients across all slices (Fig 1D, Supp. Fig 1). At baseline GCaMP activity, the mean LFP line-length and MUA firing rate were 147.9 +/- 9.2 µV/s and 4.2 +/- 0.6 spikes/s, respectively, while during GCaMP transients, these values significantly increased to 5,341.1 +/- 664.1 µV/s and 36.9 +/- 1.7 spikes/s (Fig 1E). These findings confirmed that in this slice model, GCaMP transients represented population-level excitation, as they coincided both spatially and temporally with large amplitude changes in LFP and bursts of MUA.

**Figure 1.**
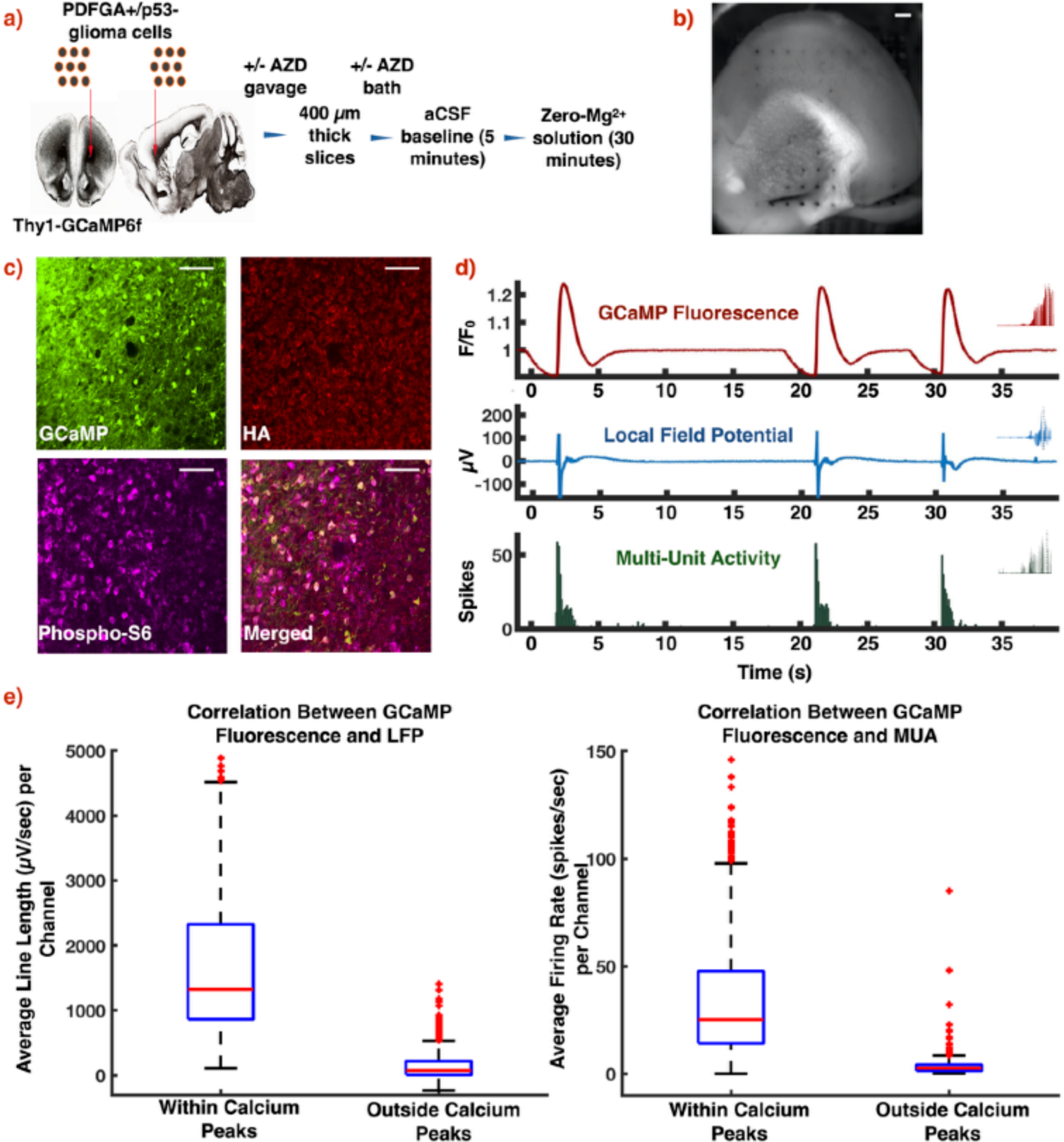
Model of diffusely infiltrating glioma and experimental design. (a) PDGFA+/p53-glioma cells were injected into the subcortical white matter of Thy1-GCaMP6f mice. Coronal slices were obtained 25-30 days post-injection. A subset of mice was treated with a single dose of AZD8055 prior to injection, and slices from these mice were additionally bathed in AZD8055 during incubation. Two slices from each mouse underwent recording in aCSF and zero-Mg2+. (b) Orientation of slice on the array in preparation for optical mapping of Thy1-GCaMP fluorescence. (c) Immunofluorescence micrographs demonstrating Thy1-GCaMP positive neurons and HA positive glioma cells. mTOR signaling was present in both Thy1-GCaMP positive neurons and HA positive glioma cells, as indicated by the presence of phosphorylated S6 ribosomal protein. (d) Representative GCaMP fluorescence, LFP, and MUA from the same channel in a tumor-bearing slice. (e) Boxplots showing temporal correlation between GCaMP fluorescence with LFP (n = 621, p < 0.01, Mann-Whitney U test, two-tailed) and MUA (n = 621, p < 0.01, Mann-Whitney U test, two-tailed). Scale bars: (b) 400 µm, (c) 50 µm.

### Tumor-bearing slices displayed increased excitability compared to age-matched controls at a population level

Previously published *in vivo* data from our Thy1-GCaMP6f murine glioma model showed the development of frequent interictal events and seizures, which appeared as high amplitude GCaMP fluctuations most prominent at the margins of the tumor^13^. By combining widefield optical imaging with high density MEA recordings, we were able to demonstrate increased excitability in acute *ex vivo* slices from these mice. We measured local responses of Thy1-GCaMP pyramidal cells to electrical stimulation in the presence of the GABA_A_ receptor antagonist picrotoxin, which blocked inhibitory synaptic transmission and facilitated assessment of excitability (Supplementary Movie 2). The mean GCaMP peak amplitude (F/F_0_) within a 140 µm radius of electrodes (Methods) which demonstrated a response at 10 µA, 25 µA, and 50 µA was 1.55 +/- 0.04, 1.58 +/- 0.03, and 1.54 +/- 0.03 in tumor-bearing slices and 1.33 +/- 0.03, 1.36 +/- 0.02, and 1.30 +/- 0.02 in controls (Fig 2B).

**Figure 2.**
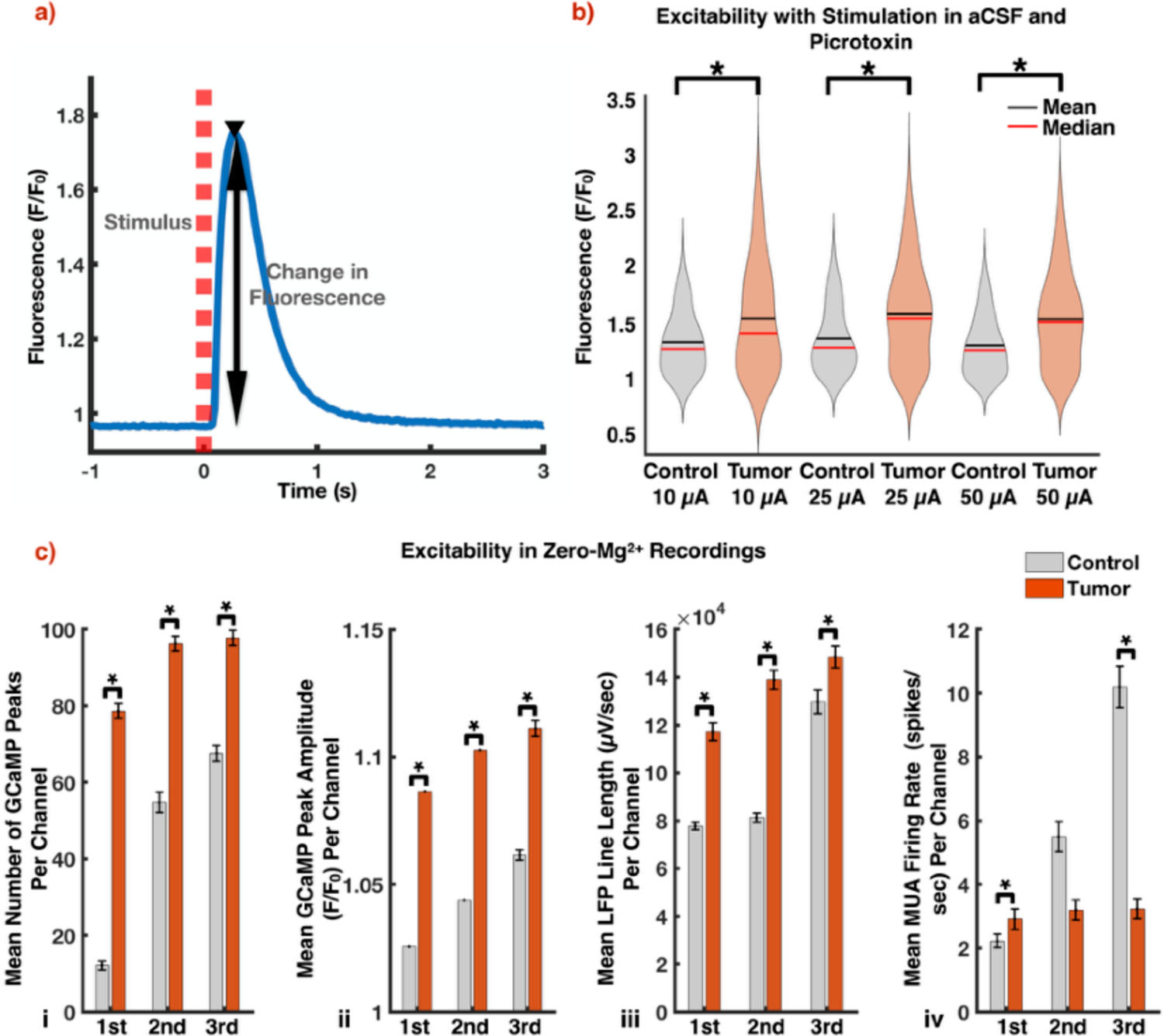
Tumor-bearing slices exhibit increased excitability compared to controls. (a) Representative local Thy1-GCaMP response (solid blue line) to microelectrode stimulation (dashed red line). (b) Violin plots displaying increased excitability in tumor-bearing slices. The mean GCaMP peak amplitude (F/F0) at electrodes which demonstrated a response at 10 µA, 25 µA, and 50 µA was 1.55 +/- 0.04, 1.58 +/- 0.03, and 1.54 +/- 0.03 in tumor-bearing slices and 1.33 +/- 0.03, 1.36 +/- 0.02, and 1.30 +/- 0.02 in controls (10 µA control: n = 79,10 µA tumor: n = 135, 25 µA control: n = 226, 25 µA tumor: n = 274, 50 µA control: n = 255, 50 µA tumor: n = 280, p < 0.01 in each, Mann-Whitney U test, two-tailed). (c) Tumor-bearing slices were more excitable than controls with respect to mean number of GCaMP peaks per channel (i), mean GCaMP peak amplitude per channel (ii), and mean LFP line length per channel (iii) (control: n = 480, tumor: n = 883, p < 0.01 for each analysis in each period, Mann-Whitney U test, two-tailed). The MUA firing rates (control: n = 480, tumor: n = 883, p < 0.01 in first period, p = 0.67 in second period, p < 0.01 in third period, Mann-Whitney U test, two-tailed) (iv) were discordant with these results.

Increased excitability in tumor-bearing slices was also displayed in zero-Mg^2+^ recordings, which were divided into three 10-minute periods. The mean number of GCaMP peaks per channel, mean GCaMP peak amplitude (F/F_0_) per channel, and mean LFP line length per channel were significantly increased in tumor-bearing slices throughout all three periods (Fig 2C). Since GCaMP and LFP reflect pyramidal cell and local summed synaptic activity^26^, respectively, these findings recapitulated increased excitability in tumor-bearing slices. The MUA firing rates, however, were greater in the control slices during the second and third periods (Fig 2C).

Considering that the MUA reflects the collective firing of all nearby neurons, including both excitatory and inhibitory ones^27^, our population-level analysis required further insight.

### Single unit analysis demonstrates entrainment of putative excitatory and inhibitory firing in tumor-bearing slices

We performed single-unit analysis to explore the behavior of individual neurons from both excitatory and inhibitory neuronal populations. 877 total units were sorted from zero-Mg^2+^ recordings in tumor-bearing slices versus 858 in controls. By using standard methods, each unit was sub-classified as a regular-spiking (RS) cell, which is largely representative of putative pyramidal neurons, or a fast-spiking (FS) cell, which is largely representative of putative PV inhibitory interneurons, by their action potential waveforms^28, 29^ (Methods). The waveform trough-to-peak and half-width durations exhibited two automatically detected well-defined clusters (Fig 3A, B). Correlating the instantaneous firing rate of each unit to the local GCaMP amplitude over time independently confirmed this subclassification across both control and tumor-bearing slices, with a higher mean correlation coefficient for RS cells than for FS cells (0.21 +/- 0.01 and 0.08 +/- 0.01, respectively; FS: *n* = 258, RS: *n =* 1000, p < 0.01, Mann-Whitney U test, two-tailed).

**Figure 3.**
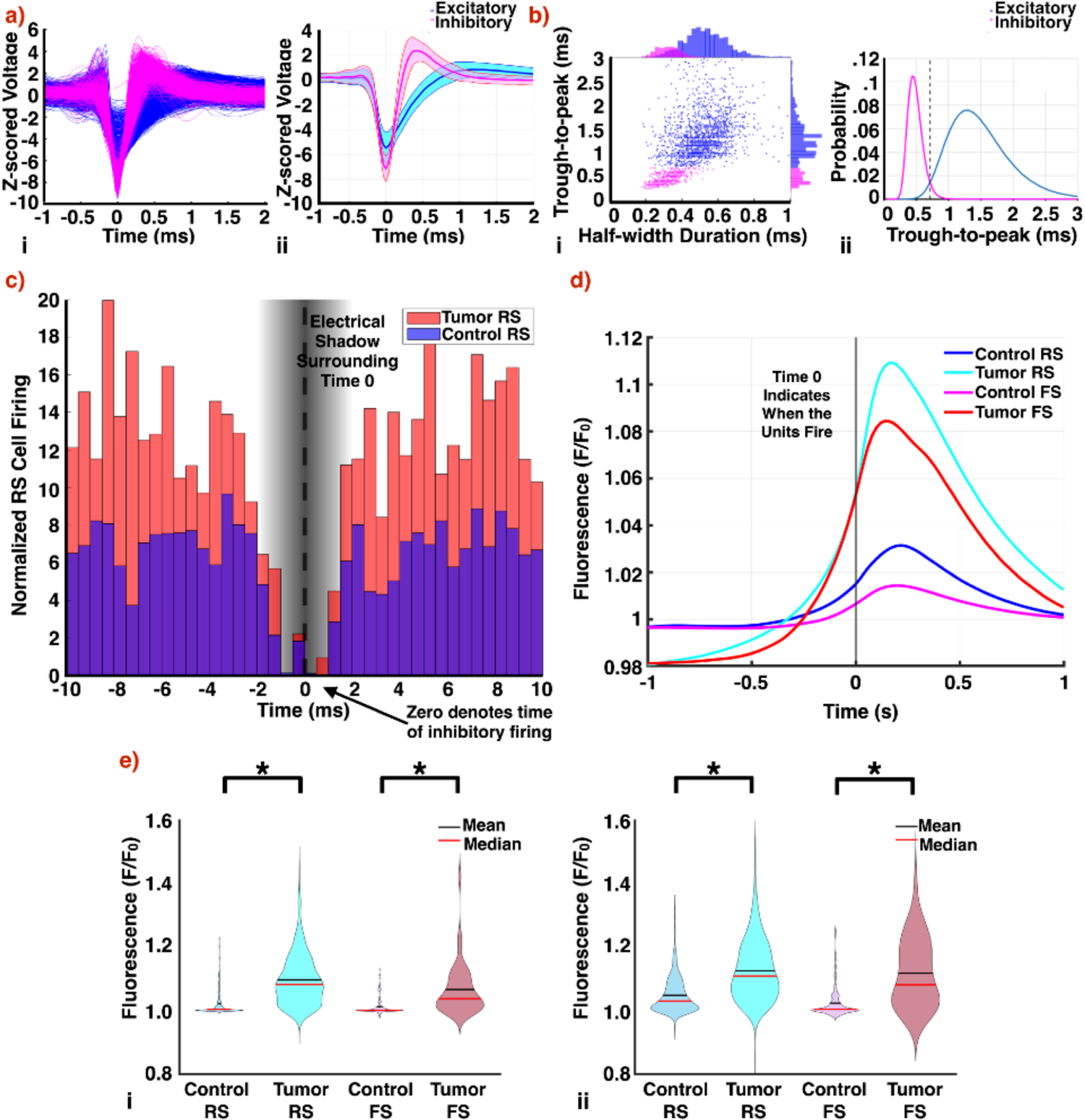
Single unit analysis demonstrates decreased inhibitory detection with dysfunctional inhibitory firing in peritumoral regions. (a) Waveforms of all putative RS cells (blue) and FS cells (magenta) (i) after sub-classification with average waveform (ii) of both groups. Shading represents SD. (b) Trough-to-peak and half-width maximum were the two parameters chosen to describe spike waveforms. Each cell’s average waveform is represented in the 2D space of the two parameters with outliers excluded (i). A bimodal distribution was evident in the trough-to-peak data, and a Gaussian mixture model was fitted to find the intersection point (0.7165 ms, dashed line) between the two components (ii). (c) Average of all cross-correlograms constructed for pairs of RS cells and FS cells sorted from the same channel. Time 0 indicates FS cell firing, while bars represent RS cell firing from −10 to 10 ms. Because of the nature of spike detection, the values of the cross-correlograms from −1.5 to 1.5 ms are thus artificially reduced. While the firing rates of FS cells were decreased in tumor-bearing slices, their activity was more concentrated within periods of increased pyramidal cell firing (control: n = 602 pairs, tumor: n = 92 pairs, p < 0.01, Mann-Whitney U test, two-tailed). (d) Mean STA GCaMP signals of each cell type in tumor-bearing slices and controls. Time 0 indicates when units fired. (e) Violin plots depicting the maximum amplitude of the STA GCaMP signals from 0 to 200 ms by cell type. In the first half of the recording (i), the median RS cell STA GCaMP signal amplitude (F/F0) was 1.08 in tumor-bearing slices versus 1.01 in controls (control: n = 350, tumor: n = 768, p < 0.01, Mann-Whitney U test, two-tailed); the median FS cell STA GCaMP signal amplitude (F/F0) was 1.04 in tumor-bearing slices versus 1.00 in controls (control: n = 204, tumor: n = 41, p < 0.01, Mann-Whitney U test, two-tailed). In the second half of the recording (ii), the median RS cell STA GCaMP signal amplitude (F/F0) was 1.08 in tumor-bearing slices versus 1.03 in controls (control: n = 376, tumor: n = 784, p < 0.01, Mann-Whitney U test, two-tailed); the median FS cell STA GCaMP signal amplitude (F/F0) was 1.11 in tumor-bearing slices versus 1.00 in controls (control: n = 204, tumor: n = 43, p < 0.01, Mann-Whitney U test, two-tailed).

Five percent (n = 44) of units from tumor-bearing slices were subclassified as FS cells, with the remaining 95% (n = 833) subclassified as RS cells. In comparison, 31% (n = 268) of units in controls were subclassified as FS cells, with the remaining 69% (n = 590) subclassified as RS cells. The difference in FS cell detection rates was likely due to a combination of both structural and functional factors, namely glioma-driven PV interneuron loss^3, 7^ and decreased PV interneuron firing rates^3^. While there was no difference in excitatory RS cell firing rates between the two cohorts (0.23 +/- 0.01 spikes/s in tumor vs 0.26 +/- 0.02 spikes/s in control, tumor: *n* = 822, control: *n* = 590, p = 0.85, Mann-Whitney U test, two-tailed), the FS cells fired slower in tumor-bearing slices at 0.26 +/- 0.03 spikes/s versus 0.65 +/- 0.05 spikes/s in controls (control: *n* = 268, tumor: *n* = 44, p < 0.01, Mann-Whitney U test, two-tailed). The decreased detection of inhibitory units and inhibitory firing in peritumoral regions both contributed to the overall decreased multiunit firing seen in tumor-bearing slices.

We constructed normalized pairwise cross-correlograms to characterize the temporal relationship between the firing of RS cells and FS cells sorted from the same electrode (602 pairs in control, 92 pairs in tumor) (Fig 3C). Cross-correlograms were normalized to the firing rates of both the FS cell and RS cell in each pair, to account for baseline firing differences by cohort (Methods).

This allowed us to analyze the response of FS cells to RS cell bursting induced by zero-Mg^2+^ solution (Fig 3C). While the firing rates of FS cells were decreased in tumor-bearing slices, their activity was more concentrated within periods of increased RS cell firing (Fig 3C). In control slices, however, FS cell firing was also maintained between bursts. Spike-triggered averaging (STA) of the time-locked local GCaMP signal to single unit firing corroborated the relationship between FS cell firing and excitatory activity in both cohorts (Fig 3D). The maximum amplitude of the STA GCaMP signal from 0 to 200 ms was recorded for both RS cells and FS cells. These values were significantly higher in tumor-bearing slices for FS cells across the entire recording (Fig 3E). This highlights how FS cell firing in tumor-bearing slices is associated with increased pyramidal activity.

### mTOR inhibition reduces peritumoral excitability

The mTOR pathway is of special interest as its upregulation has been demonstrated in both glioma^30, 31^ and epilepsy^32^. We thus used AZD8055 (AZD), a competitive inhibitor of the kinase domains of mTORC1 and mTORC2, to characterize the functional consequences of mTOR inhibition on specific neuronal populations within the peritumoral microenvironment. A single oral gavage of AZD (100 mg/kg) was administered to tumor-bearing mice 4-5 hours prior to sacrifice. Slices obtained from these mice were additionally bathed in 30 nM of AZD during their incubation period prior to recording. Immunofluorescence analysis demonstrated that the single dose of AZD significantly reduced levels of phospho-S6 in both neuronal and neoplastic cells across the slice (Fig 4A-C).

**Figure 4.**
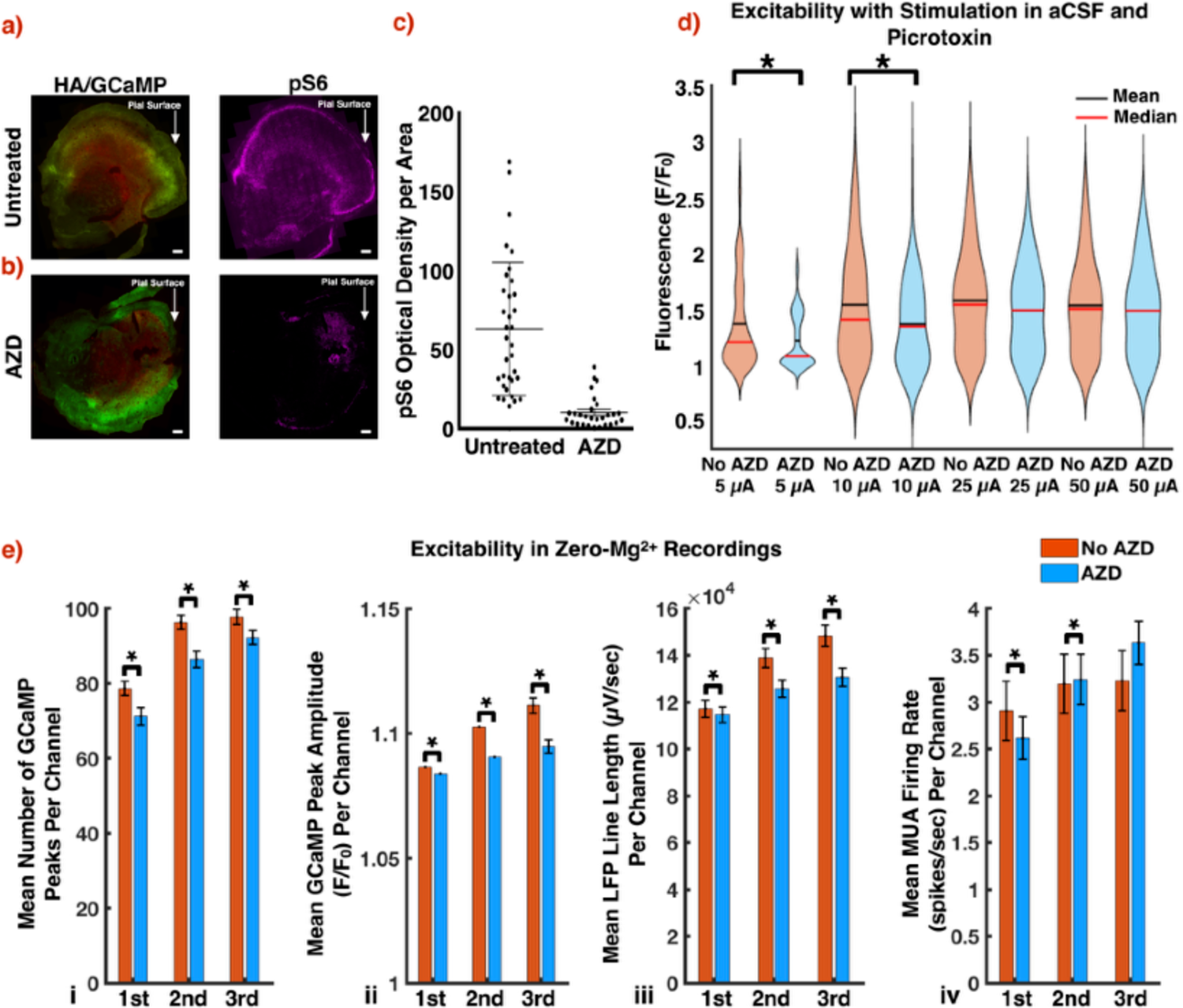
mTOR inhibition decreases peritumoral excitability. Low power immunofluorescence micrographs of untreated. (a) and treated (b) tumor-bearing slices, demonstrating decreased mTOR signaling in neuronal and neoplastic cells of treated slices. This was evidenced by reduced levels of phosphorylated S6 ribosomal protein (untreated: n = 34, treated: n = 27, p < 0.01, unpaired t-test, two-tailed) (c). (d) Violin plots displaying decreased excitability in treated tumor-bearing slices. The mean amplitudes of the local GCaMP response (F/F0) at 5 µA was 1.37 +/- 0.04 in untreated slices versus 1.23 +/- 0.03 in treated ones (5 µA untreated tumor: n = 102, 5 µA treated tumor: n = 53, p < 0.05, Mann-Whitney U test, two-tailed); at 10 µA it was 1.55 +/- 0.04 in untreated slices versus 1.37 +/- 0.04 in treated ones (10 µA untreated tumor: n = 135, 10 µA treated tumor: n = 116, p < 0.01, Mann-Whitney U test, two-tailed). (e) Excitability was decreased in treated tumor-bearing slices with respect to mean number of GCaMP peaks per channel (i), mean GCaMP peak amplitude per channel (ii), and mean LFP line length per channel (iii) (untreated tumor: n = 883, treated tumor: n = 916, p < 0.01 for each analysis in each period, Mann-Whitney U test, two-tailed). The MUA firing rates (untreated tumor: n = 883, treated tumor: n = 916, p < 0.05 in first period, p < 0.01 in second period, p = 0.12 in third period, Mann-Whitney U test, two-tailed) (iv) were discordant with these results. Scale bars: (a) 400 µm, (b) 400 µm.

We reassessed peritumoral excitability in the setting of mTOR inhibition. In comparison to untreated tumor-bearing slices, direct stimulation of treated ones in the setting of GABAergic antagonism with picrotoxin displayed decreased mean amplitudes of the local GCaMP response at 5 µA (1.37 +/- 0.04 versus 1.23 +/- 0.03 F/F_0_) and 10 µA (1.55 +/- 0.04 versus 1.37 +/- 0.04 F/F_0_) (Fig 4D).

Decreased excitability in treated tumor-bearing slices was also demonstrated in zero-Mg^2+^ recordings (Fig 4E). The mean number of GCaMP peaks per channel, mean GCaMP peak amplitude (F/F_0_) per channel, and mean LFP line length (µV/s) per channel were significantly decreased in treated slices throughout all three periods of the recording (Fig 4E). While these three metrics showed decreased peritumoral excitability after mTOR inhibition, we were unable to draw similar conclusions from an assessment of the multiunit firing rates in these slices (Fig 4E). This called for an evaluation of mTOR inhibition’s effects on individual neuronal populations.

### Peritumoral interneurons demonstrate decreased entrainment to RS cell firing after mTOR inhibition

Single unit analysis was performed for treated tumor-bearing slices as described above. 794 total units were sorted from treated slices, of which 15% (n = 119) were sub-classified as FS cells, an increase from the 5% detected in untreated tumor-bearing slices (Fig 5A). Treated FS cells fired at an average rate of 0.70 +/- 0.09 spikes/s, exceeding the 0.26 +/- 0.03 spikes/s seen in untreated tumor-bearing slices (Fig 5B). There was no significant difference in RS cell firing rates.

**Figure 5.**
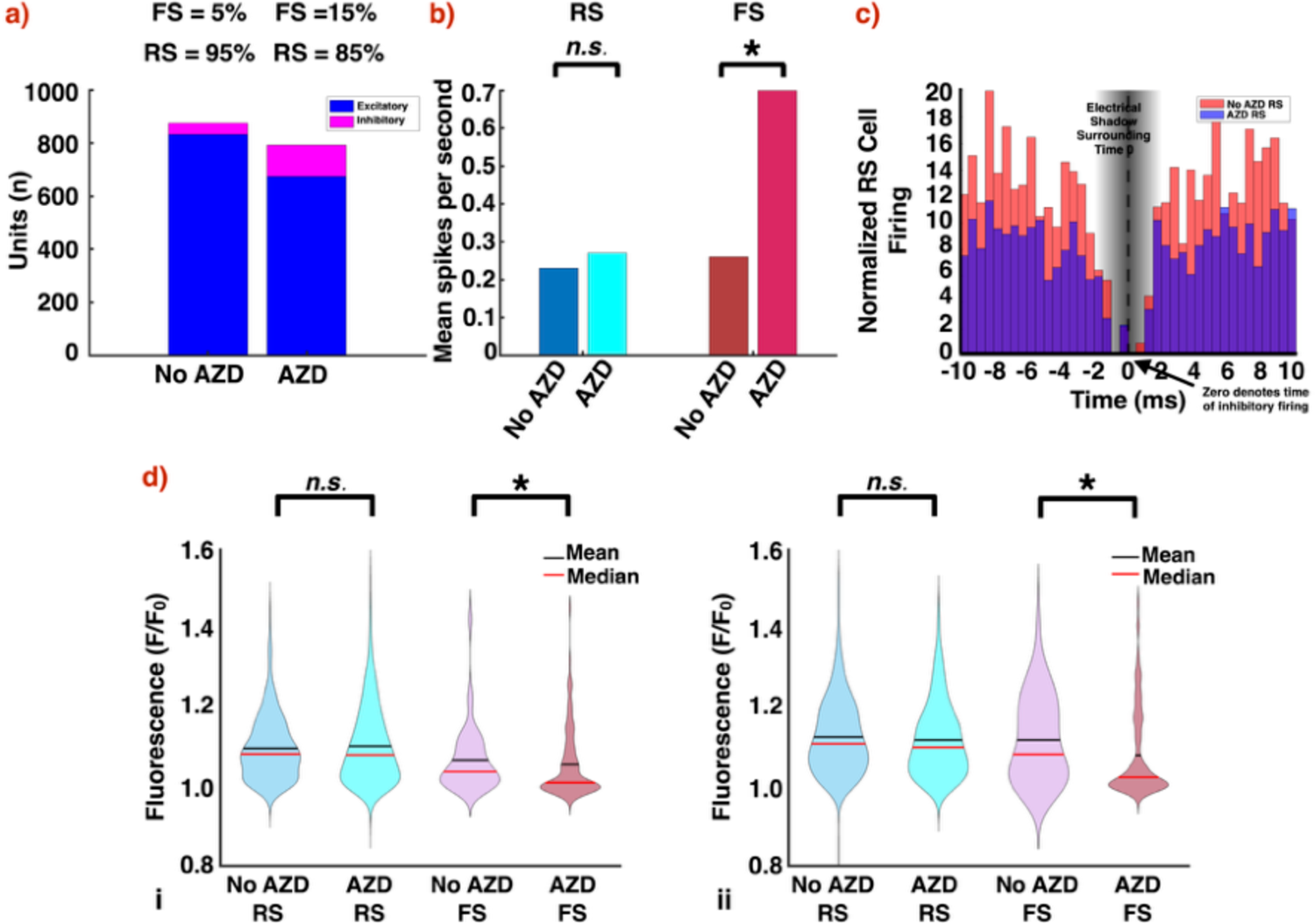
Peritumoral FS cells demonstrate decreased entrainment to RS cell firing after mTOR inhibition. (a) In treated tumor-bearing slices, 15% of sorted units were sub-classified as FS cells versus 5% in untreated slices. (b) FS cells from treated slices fired at 0.70 +/- 0.09 spikes/s, as opposed to 0.26 +/- 0.03 spikes/s in untreated slices (untreated tumor: n = 44, treated tumor: n = 119, p < 0.01, Mann-Whitney U test, two-tailed). (c) In comparison to those from untreated tumor-bearing slices, pairwise cross-correlograms from treated ones revealed decreased RS cell firing in temporal relationship to FS cell firing, suggesting greater interictal FS cell activity (untreated tumor: n = 92 pairs, treated tumor: n = 199 pairs, p < 0.01, Mann Whitney U test, two-tailed). (d) Similarly, spike-triggered averaging of the local GCaMP response to FS cell firing was decreased in treated tumor-bearing slices in the first (untreated tumor: n = 41, treated tumor: n = 117, p < 0.05, Mann-Whitney U test, two-tailed) (i) and second (untreated tumor: n = 43, treated tumor: n = 118, p < 0.01, Mann-Whitney U test, two-tailed) (ii) halves of the recording. There was no significant difference for spike-triggered averaging of the local GCaMP responses to RS cell firing.

In comparison to those from untreated tumor-bearing slices, pairwise cross-correlograms (199 pairs) from treated slices revealed decreased RS cell firing in temporal relationship to FS cell firing (Fig 5C). Similarly, STAs of the local GCaMP signal to FS cell firing were significantly decreased in treated tumor-bearing slices (Fig 5D). There was no difference between STAs of the local GCaMP signal to RS cell firing. These findings suggested increased FS cell firing between excitatory bursts in treated tumor-bearing slices.

### mTOR inhibition restores FS cell-mediated surround inhibition during ictal events

Ictal events are characterized by a spatial restraint in which inhibitory barrages from FS cells prevent the propagation of epileptiform activity^33–39^. Prior studies have shown that interneurons located outside the region bounded by the ictal wavefront contribute to surround inhibition^36^. In contrast, interneurons focal to the ictal activity have been implicated in its propagation^34^ and have been shown to alter their behavior prior to and during seizures^7, 40, 41^. These interneuron states, however, have yet to be characterized in the setting of TAS. Consequently, we sought to assess how the spatial landscape of surround inhibition was impacted by our findings of glioma-induced FS cell dysfunction.

We combined widefield optical imaging with single unit analysis to characterize the spatial relationship between FS cell firing and propagating GCaMP transients. For each FS cell, we obtained the Euclidean distance between the electrode at which the FS cell firing was detected, and every other electrode on the array. We then calculated the maxima of the average time-locked GCaMP signal at each electrode during a 400 ms window around the cell’s firing. Across all FS cells, the maxima at each unique distance were averaged together to determine how GCaMP activity varied with distance from FS cell firing (Methods).

With this technique, we were able to identify FS cells both focal to and distant from excitatory activity in our control slices (Fig 6A). We observed increased levels of GCaMP activity both proximal to and distant from regions with FS cell firing. This was visualized as a bimodal distribution when plotting the average maximum fluorescence at each unique distance across all FS cells (Fig 6A). We interpreted this as evidence of interneurons which have either been recruited to or lie outside of epileptiform bursts, with the latter suggesting surround inhibition. To deconvolve these two subpopulations, we evaluated the maxima of the average time-locked GCaMP signal at each unique distance for each FS cell. If the signal peaked within 0.4 mm (one electrode) of the FS cell, that cell was classified as recruited to excitatory activity (Methods). Otherwise, the cell was classified as contributing to surround inhibition (Methods).

**Figure 6.**
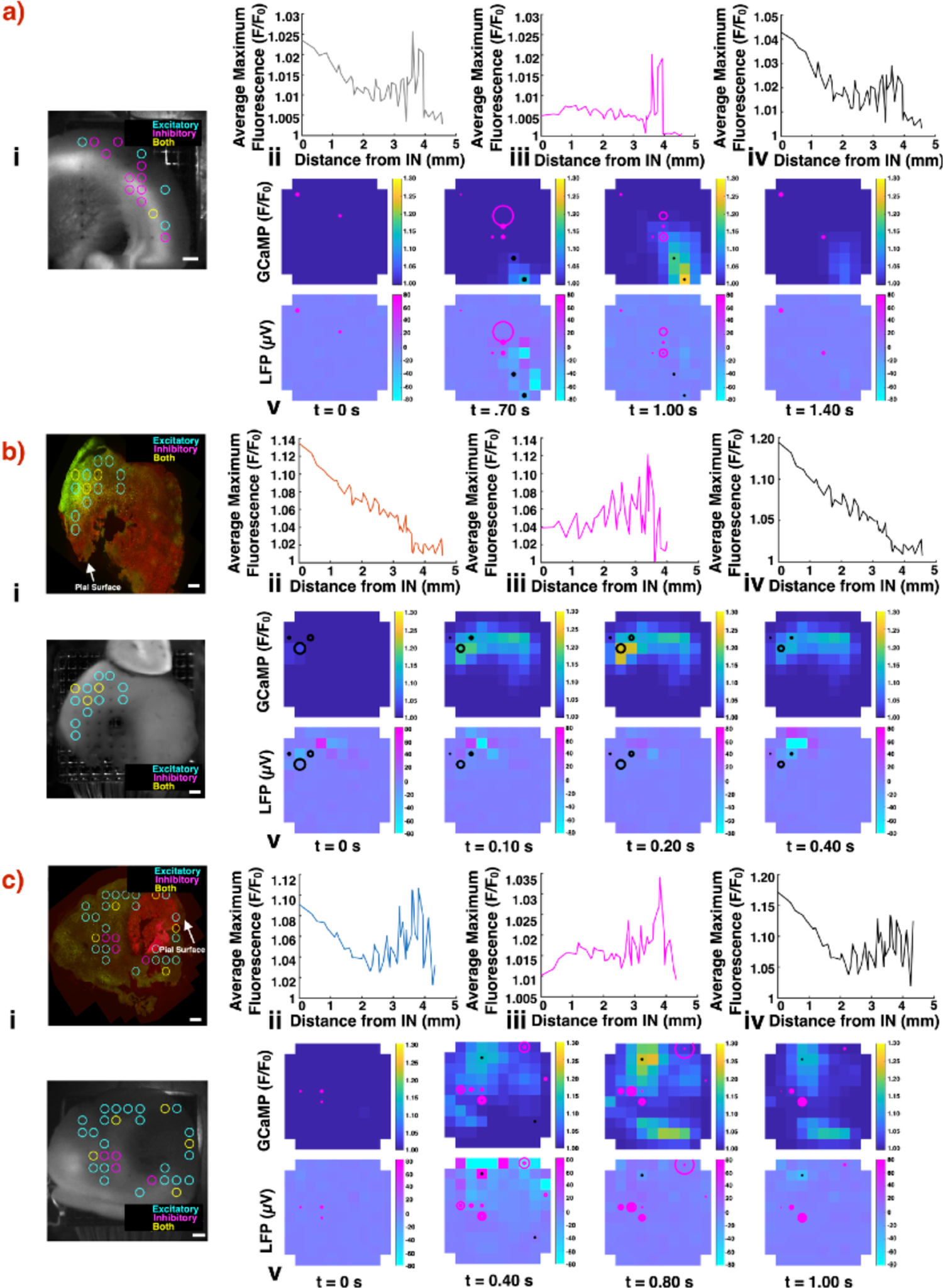
mTOR inhibition restores FS cell-mediated surround inhibition during ictal events. (a) In control slices, the time-locked GCaMP signal around FS cell firing suggested intact surround inhibition. Areas with increased GCaMP activity were located both proximal to and distal from areas with FS cell firing (ii). Four consecutive time points are shown (v) from Supplementary Movie 3 during a representative ictal event from a control slice. The representative slice shows that maximum GCaMP activity occur at a distance from the FS cells. Moreover, the FS cells appear to inhibit spread. (b) In untreated tumor-bearing slices, the time-locked GCaMP signals with the greatest amplitudes were proximal to electrodes with FS cell firing (ii). As distance from the electrodes with these FS cells increased, the time-locked GCaMP signal amplitude decreased. Four consecutive time points are shown (v) from Supplementary Movie 4 during a representative ictal event from an untreated tumor-bearing slice. The representative slice shows maximum GCaMP activity occurring in the electrodes with FS cell firing. (c) Treated tumor-bearing slices recapitulated the findings seen in controls, suggesting that mTOR inhibition restores the contribution of FS cells to surround inhibition during ictal events (ii). Four consecutive time points are shown (v) from Supplementary Movie 5 during a representative ictal event from a treated tumor-bearing slice. The representative slice shows maximum GCaMP activity occurring at a distance from FS cell firing. Across all cohorts, areas of increased GCaMP activity occurred distal to FS cells engaged in surround inhibition (iii) and proximal to FS cells recruited to excitatory activity (iv) (a-c). Control and treated tumor-bearing slices possessed a greater percentage of FS cells contributing to surround inhibition (50.7% and 50%, respectively) compared to untreated tumor-bearing slices (9.3%). Representative histology (HA positive glioma cells, red; Thy1 positive neurons, green) shown with pictures of slices placed on the array (i). Scale bars: 400 µm. Heatmaps denote GCaMP (top) and LFP (bottom) amplitude. Radii of circles (magenta for FS cells contributing to surround inhibition, black for FS cells recruited to excitatory activity) on heatmaps indicate the instantaneous firing rate of the FS cells in that electrode.

In control slices, 104 FS cells (50.7%) were classified as contributing to surround inhibition, while 101 (49.3%) were classified as recruited (Fig 6A). The average firing rate of FS cells contributing to surround inhibition was 0.95 +/- 0.10 spikes/s, and the average firing rate of FS cells recruited to excitatory activity was 0.66 +/- 0.08 spikes/s (surround inhibition: *n* = 104, recruited to excitatory activity: *n* = 101, p < 0.01, Mann-Whitney U test, two-tailed). The observed reduction in the firing rate of the recruited FS cells suggests a breakdown in inhibitory firing. Supplementary Movie 3 displays the spread of excitatory activity (GCaMP, LFP) across a representative control slice, with FS cell firing overlays (magenta circles for FS cells contributing to surround inhibition and black circles for FS cells recruited to excitatory activity, whose radii are determined by their instantaneous firing rates relative to elsewhere in the recording). The four consecutive time points shown demonstrate the presence of FS cells both recruited to areas of excitatory activity or contributing to surround inhibition, illustrating how the observed epileptiform burst did not extend beyond the region containing the latter (Fig 6A).

In untreated tumor-bearing slices, maximum GCaMP activity was observed more within areas of FS cell firing and less in regions distant to the interneurons. As distance from the FS cells increased, the average maximum fluorescence decreased, suggesting a loss of inhibition (Fig 6B). Thirty-nine FS cells (90.1%) were classified as recruited to excitatory activity, while only 4 (9.9%) were classified as contributing to surround inhibition. There was no significant difference in the firing rates between these two groups. A representative ictal event from an untreated tumor-bearing slice can be seen in Supplementary Movie 4. The four consecutive time points shown highlight how the epileptiform activity begins in the region with FS cell firing, which we interpreted as indicative of a loss of surround inhibition (Fig 6B).

Treated tumor-bearing slices demonstrated findings similar to that of control slices. Increased levels of GCaMP were observed both proximal to and distant from regions with FS cell firing (Fig 6C). Fifty-nine FS cells (50.0%) were classified as contributing to surround inhibition, which was increased from the 9.9% seen in untreated tumor-bearing slices. The average firing rate of FS cells contributing to surround inhibition was 1.03 +/- 0.16 spikes/s, and the average firing rate of FS cells recruited to excitatory activity was 0.38 +/- 0.08 spikes/s (surround inhibition: *n* = 59, recruited to excitatory activity: *n* = 59, p < 0.01, Mann-Whitney U test, two-tailed). Supplementary Movie 5 showcases epileptiform activity in a treated tumor-bearing slice. The four consecutive time points highlight how regions with maximum GCaMP fluorescence are at a distance from FS cells contributing to surround inhibition (Fig 6C). These findings are consistent with a restoration of surround inhibition within the peritumoral cortex following treatment with AZD.

## Discussion

We combined optical mapping of Thy1-GCaMP6f fluorescence with single unit analysis from multielectrode extracellular recordings, acquiring both population and individual-level cell-type specific data across an extended peritumoral cortical region. This allowed us to interrogate the spatiotemporal behavior of both putative pyramidal cells and PV interneurons in a diffusely infiltrating model of glioma. Moreover, this approach provided an electrophysiologic correlate for the GCaMP fluctuations observed in longitudinal *in vivo* imaging of this model^13^. In agreement with prior studies, our *ex vivo* acute tumor-bearing slices demonstrated increased excitability^4, 7, 12, 17, 42^ and decreased FS cell firing^3^ without significant increases in pyramidal cell activity. Additionally, peritumoral FS cell firing was less likely to be interictal, as it occurred predominantly within epileptiform bursts. This breakdown in inhibitory firing was associated with deficits in surround inhibition. These findings suggest that increased excitability is correlated with inhibitory dysfunction, building upon reports of GABAergic disinhibition in TAS^2, 7^. In contrast with existing hypotheses regarding structural mechanisms of inhibitory dysfunction, the glioma-induced alterations to the spatiotemporal relationship between excitatory and inhibitory neurons were reversed by inhibiting mTOR.

Increased mTOR signaling has been observed in both the tumor core and the non-neoplastic cells of the peritumoral microenvironment^31, 43^. mTOR hyperactivation within neurons in particular can lead to dysfunctional axon production, enlarged neuronal soma size, upregulation of immature N-Methyl-D-aspartic acid (NMDA) receptors^25^, and decreased GABAA-receptor trafficking^44^. All of these changes can contribute to increased excitability. In this study, we perturbed our *ex vivo* acute slice model of diffusely infiltrating glioma with the mTOR inhibitor AZD, which inhibits the kinase domains of both mTORC1 and mTORC2^45^. This treatment reduced excitability, as evidenced by both local stimuli delivered in picrotoxin and recordings performed in zero-Mg^2+^ solution. These results highlight a role for mTOR signaling in alterations of the excitatory-inhibitory balance at the infiltrative margins of glioma.

Glioma-induced changes to PV interneurons have been implicated in the generation of TAS and likely account for the alterations in FS cells seen in our study. Within the peritumoral microenvironment, PV interneurons are decreased in number^2, 3, 7^ and fire at slower rates^3^.

Moreover, glioma-induced changes to chloride regulation in pyramidal neurons result in paradoxical GABAergic excitatory responses^2, 7^. Using our model, we show that peritumoral FS cells and their effects on the excitatory-inhibitory landscape can be modulated through mTOR inhibition. In treated tumor-bearing slices, FS cell firing was increased but less concentrated around pyramidal bursts, suggesting greater interictal FS cell firing. This result provides evidence that glioma-induced changes to the excitatory-inhibitory balance extend beyond structural ones^3, 7^. Indeed, we show that some are functional and partially reversible.

Previous *ex vivo*^33, 35, 36^, *in vivo*^37^, and human^46^ studies have demonstrated the contribution of PV interneurons, of which FS cells are representative^28, 29^, to surround inhibition during ictal events. In such instances, PV interneurons rapidly respond to excitatory firing and restrain spatial propagation^33–38^. Functional impairment of these interneurons leads to collapse of local inhibition and facilitates seizure spread^35^. Herein, we combined single unit analysis with cell type-specific GCaMP imaging to characterize surround inhibition in acute *ex vivo* slices. This allowed us to spatially investigate whether putative PV interneurons were contributing to inhibition or recruited to ongoing ictal activity. Both states were seen in control slices, where increased GCaMP fluorescence was observed distal to some FS cells and proximal to others. In tumor-bearing slices, increased GCaMP fluorescence was found on average only proximal to FS cells.

We attributed this to a glioma-induced breakdown in surround inhibition. Tumor-bearing slices treated with AZD, however, demonstrated findings similar to that of controls. This suggests that mTOR inhibition can restore FS cell-mediated surround inhibition. These findings may have implications for the clinical course of patients with diffusely infiltrating glioma. Recent reports have shown that aberrant neuronal activity can lead to glioma proliferation through various mechanisms^4, 8–10, 13, 15^. It is thus conceivable that mTOR-mediated restoration of surround inhibition could potentially reduce the spread of such aberrant firing and in turn decrease tumor progression. However, we acknowledge that this analysis was limited by a lack of direct imaging of PV interneurons and an investigation of shorter ranges of inhibition, calling for additional studies.

In conclusion, we have described the spatiotemporal behavior of individual neuronal populations in an acute *ex vivo* model of diffusely infiltrating glioma. By doing so, we have demonstrated alterations in the excitatory-inhibitory landscape of the peritumoral cortex, including increased excitability, a reduction in FS cell interictal firing, and a loss of surround inhibition. These changes were partially reversed with a single dose of AZD, suggesting that functional mechanisms linked to mTOR activation are prominently involved. Our findings have implications for both seizure susceptibility and glioma progression, warranting further analysis of mTOR’s role in tumor-associated seizures.

## Methods

### Murine glioma model

All animal handling and experimentation were performed with the approval of the Columbia University Institutional Animal Care and Use Committee. As previously described, diffusely infiltrating glioma were generated by injecting PDGFA-IRES-Cre expressing retrovirus into the subcortical white matter of transgenic C57BL/6 mice that possessed floxed p53, stop-flox RPL22^HA^, and stop-flox mCherry-luciferase^13^. The resulting retrovirus-induced tumors show the histological features of diffusely infiltrating gliomas^13^. Glioma cells were then isolated from these retrovirally-induced tumors and sustained in culture^47^. These cells were injected into the subcortical white matter of adult (∼2 months old) Thy1-GCaMP6f positive mice of both sexes (stereotaxic coordinates relative to the bregma: 2.0 mm lateral, 2.0 mm rostral, and 1.5 mm deep)^13^. The mice readily formed tumors within 2-3 weeks and developed spontaneous behavioral seizures as early as 3-weeks post-injection^13^.

### Slice preparation

Acute brain slices were prepared from tumor-bearing mice (n = 12) sacrificed 25 to 30 days post-injection; slices were also acquired from non-injected age-matched controls (n = 3). 400 µm thick coronal slices were obtained using a Leica VT1000S vibratome (Nussloch, Germany) while in ice-cold solution [in mM: 210 Sucrose, 10 Glucose, 2.5 KCl, 0.5 CaCl_2_, 7 MgCl_2_, 26 NaHCO_3_, 1.25 NaH_2_PO_4_]. 4 slices were obtained from each mouse. The slices were incubated at 35 °C for 18 min and then transferred to 22 °C for a minimum of 42 min prior to recordings.

Contents of the incubation solution are listed as follows [in mM: 1.5 MgCl_2_, 125 NaCl, 26 NaHCO_3_, 10 dextrose, 2.5 KCl, 2 CaCl_2_, 1.25 NaH_2_PO_4_].

### AZD8055 treatment

Half of the tumor-bearing mice (n = 6) underwent a single oral gavage with 100 mg/kg of AZD, a competitive inhibitor of the kinase domains in mammalian target of rapamycin complex 1 (mTORC1) and mammalian target of rapamycin complex 2 (mTORC2), 4-5 hours prior to sacrifice. This time window was selected as we have shown that oral administration of AZD decreases neuronal mTOR activation *in vivo* within 6 hours of administration as evidenced by decreased phosphorylation of S6 (D. Torres, PC, unpublished data). The slices obtained from these mice were additionally bathed in 30 nM of AZD during their incubation period prior to recording. The remaining tumor-bearing mice (n = 6) and non-injected age-matched controls remained untreated.

### Slice recordings and wide-field calcium imaging

Slices were placed on top of a 4 × 4 mm MEA with 96 (10 × 10 electrode grid, non-recording corners) 1 mm penetrating microelectrodes at an orthogonal interelectrode spacing of 0.4 mm (Blackrock Microsystems, Inc., Salt Lake City, UT). The electrode array sampled from across the slices, including both cortical and subcortical regions.

Two slices from each mouse underwent an initial 5-minute recording in aCSF [in mM: 1.5 MgCl_2_, 125 NaCl, 26 NaHCO_3_, 10 dextrose, 5 KCl, 2 CaCl_2_, 1.25 NaH_2_PO_4_], followed by a 30-minute recording in zero-Mg^2+^. This limited our analysis to early ictal events, which better reflect clinical conditions. Signals from the MEA were acquired continuously on a CerePlex Direct acquisition system with a digital preamplifier (Blackrock Microsystems, Inc., Salt Lake City, UT) at 30 kHz per channel (0.3 Hz −7.5 kHz bandpass), with 16-bit precision and a range of +/− 8 mV.

The other two slices from each mouse underwent stimulation in aCSF with the addition of 50 µM picrotoxin, a GABA_A_ receptor antagonist. This allowed us to interrogate excitatory responses to stimulation without the presence of an inhibitory restraint. Each of the 96 electrodes were stimulated in ascending order, with odd-numbered electrodes stimulated prior to even-numbered ones. Stimulus trains consisted of 25 cathodic first symmetrical pulses with phases lasting 100 µs and separated by an interphase delay of 100 µs, delivered at a frequency of 100 Hz using the Cerestim R96 (Blackrock Microsystems Inc., Salt Lake City, UT). The total duration of each pulse train was 0.25 seconds. All electrodes were first stimulated at 5 µA, with a spacing of 3 seconds between each sequential stimulus. After each electrode had delivered this stimulus, the protocol was repeated at 10 µA, 25 µA, and 50 µA.

Imaging was performed under an upright Leica DM LFS microscope using a 2.5x/0.07NA Leica objective and an I3 fluorescence filter cube (Leica Microsystems, Wetzlar, Germany). GCaMP6f signal was excited using a wavelength band of 450 to 490 nm. Time series images were acquired at a frame rate of 50 Hz and resolution of 1024×1024 pixels using an Andor Zyla Plus sCMOS 4.2 MP camera (Oxford Instruments, Abingdon, UK).

### Histology and confocal microscopy

After recording, slices were removed from the array and fixed with 4% paraformaldehyde (PFA) overnight at 4 °C. 40 µm sections were cut with a vibratome. For immunohistochemical analysis, free-floating sections were blocked in 10% goat serum (30 minutes, room temperature) and then incubated with primary antibodies (overnight at 4 °C) for a marker of tumor cells, anti-hemagglutinin (rat monoclonal, catalog #11867423001, Millipore-Sigma, MA, RRID: AB_390918), and a marker of mTOR activation, anti-phospho-s6 ribosomal protein (pS6 Ser240/244) (rabbit monoclonal, catalog #2215S, Cell Signaling Technology, MA, RRID: AB_331682). Secondary Alexa Fluor 594 (goat anti-rabbit, catalog #A32740, Invitrogen, OR, RRID: AB_2762824) and 647 (goat anti-rat, catalog #A32733, Invitrogen, OR, RRID: AB_2633282) conjugated antibodies were then applied with DAPI for 1 hour at room temperature. Blocking serum, primary antibodies, and secondary antibodies were applied in 0.3% Triton X-100 in PBS. Thy1 positive neurons were identified using their endogenous GCaMP fluorescence.

Z-series confocal images were obtained using an inverted confocal microscope (Eclipse Ti, Nikkon, Japan) under a 20× air objective (N/A 0.75) (Nikkon, Japan) with 7 µm incremental steps. Serial images were obtained with 10% overlap and then stitched together (NIS Elements, Nikkon, Japan) to obtain an image of the whole slice. Z-series were then stacked together to generate the max-intensity projection. All confocal images shown were projected views. These images were exported and processed in Fiji^44^ and MATLAB (MathWorks, Natick, MA). Levels of pS6 fluorescence were calculated using the optical density of the slice in Fiji.

### Data processing and statistical analysis

#### LFP and MUA

For initial analyses, the raw MEA signals were symmetrically bandpass filtered into two frequency bands: LFP (2 - 50 Hz, 512th order, window-based FIR1 filter in MATLAB) and MUA (500 Hz – 5 kHz, 512th order, window-based FIR1 filter in MATLAB). Both filtered data streams were visually reviewed to exclude channels and time periods with excessive artifact.

#### Single unit discrimination

For single unit analyses, the MEA signals were symmetrically filtered between 300 Hz and 5 kHz to accommodate potential waveform changes during epileptiform activity (1024th order window-based FIR filter). Extracellular action potential waveforms were detected from this filtered signal using a threshold crossing method^48^. Putative single unit spikes were identified as voltage peaks greater than 4.5 σ where σ = median(|x|/0.6745), and x is the filtered signal from that channel. Matrices of waveforms from each channel were created from 0.6 ms prior to 1 ms post each detection, and principal component based semi-automatic cluster cutting was performed using a modified version of the “UltraMegaSort2000” MATLAB toolbox^46, 47^ and the “SplitMerge” graphical interface^18^. Clusters that satisfied the following criteria were accepted: (i) clean separation from all other clusters in the Fisher’s linear discriminant in principal component space; (ii) less than 1% contamination of the 2 ms absolute refractory period; (iii) no clear outliers based on the anticipated chi-squared distribution of Mahalanobis distances; and (iv) less than 1% of estimated false negatives as estimated by the amount of a Gaussian fit to the detected voltages fell below the threshold for detection^49^.

#### Subclassification of RS cells and FS cells

Single units were subclassified into FS cells and RS cells based on the unit’s mean waveshape (trough to peak duration and spike half-width) from the original, unfiltered signal. Waveform duration has been shown to be shorter in FS cells^50, 51^. A bimodal distribution was evident in the trough-to-peak data, and a Gaussian mixture model was fitted to find the intersection point between the two components (Fig. 3A-B). Subclassified identities were confirmed through correlation of the cell-type firing rates and local calcium imaging fluorescence.

#### Calcium imaging analysis

The average fluorescence within a 140 µm radius of each electrode^52^ was calculated for each frame of the recording. The average fluorescence in each frame was then normalized to the baseline intensity using a sliding window of 125 frames before and after, channel by channel.

### Multimodal recording analysis

#### Slices undergoing recording in zero-Mg2+

To investigate whether GCaMP fluctuations correlated with changes in electrophysiologic data, we identified calcium peaks as those 1% greater than the normalized baseline. The mean LFP line-length and MUA firing rate were then calculated within a 1 second window around each peak. The former was obtained by summing the values between each consecutive data point and dividing by one second, while the latter was acquired by dividing the total number of spikes in each window by one second. The same analysis was then performed for periods between calcium peaks. Results were pooled across all cohorts. This correlation analysis was restricted to electrodes whose overall LFP line-length and MUA firing rate were greater than the mean across the slice.

Each 30-minute recording was divided into three periods of ten minutes each to account for the evolving dynamics of activity in zero-Mg^2+^ solution. Hyperexcitability was then quantified for each period through population-level data, including Thy1-GCaMP6f fluorescence, LFP, and MUA. As above, calcium peaks were identified as those 1% greater than the normalized baseline. These were used to calculate average peak amplitude and average number of peaks per channel. Each electrode’s LFP line-length and MUA firing rate were additionally calculated as previously described.

#### Slices undergoing stimulation in aCSF + picrotoxin

The local Thy1-GCaMP response to stimulation was examined for each electrode. Amplitude was evaluated for each response. These values were recorded separately for each level of stimulation, at 5 µA, 10 µA, 25 µA, and 50 µA.

### Single unit analysis

From all sorted units, the detection rates for RS cells and FS cells were determined for each cohort. The firing rate for each unit was determined by dividing their total number of spikes by the length of the recording. Pairwise cross-correlograms were constructed for FS cell/RS cell pairs sorted from the same electrode. To account for baseline firing rate differences across slices and experimental conditions, the cross-correlograms were normalized both to the total spikes of the FS cell and to the mean firing rate of the RS cell for the given bin-width, such that a resultant value of 1 would denote the expected firing rate of the RS cell at any given time, including quiescent inter-discharge periods.

As previously described, the spike-triggered average (STA) was calculated for each individual unit, with the local Thy1-GCaMP fluorescence as the response^53^. This was done for each half of the recording. The max amplitude of the STA from 0 to 200 ms was recorded for each unit across all treatment conditions.

For each FS cell, we obtained the Euclidean distance between its electrode and every other electrode on the array. We then calculated the maxima of the average time-locked GCaMP signal at each electrode within a 400 ms window around the cell’s firing. Across all FS cells, the maxima at each unique distance were averaged together to determine how GCaMP activity varied with distance from FS cell firing. This was done separately for control, untreated tumor, and treated tumor cohorts. To differentiate between FS cells recruited to excitatory activity and those contributing to surround inhibition, we evaluated the maxima of the average time-locked GCaMP signal at each unique distance for each FS cell. If the signal peaked within 0.4 mm (one electrode) of the FS cell, that cell was classified as recruited to excitatory activity. Otherwise, the cell was classified as contributing to surround inhibition. This analysis was confined to the latter half of our recordings. This period was chosen as it was more likely to showcase epileptiform activity, enabling us to characterize the FS cell responses to it.

### Statistical analysis

All analyses were performed offline using custom scripts and toolboxes written in MATLAB (MathWorks, Natick, MA). Code is available at https://github.com/edmerix. When provided, means were reported +/- SEM. All statistical tests for significance were performed using the Mann-Whitney U test unless otherwise noted, due to the non-Gaussian distributions of data requiring non-parametric testing. For all tests, the level for statistical significance (α) was set to 0.05.

## Author Contributions

FK, BG, AG, GM, CS, and PC designed the experiments. AG and AD injected and maintained the transgenic mice. BG, FK, AG, XW and TS performed slice experiments. AS performed the immunofluorescence staining of the acute brain slices. BG, FK, AG, EM, and JL acquired, analyzed, and interpreted the data. FK and BG wrote the first version of the manuscript and all authors revised it. All authors have read and approved the final version of this manuscript. All authors agree to be accountable for all aspects of the work in ensuring that questions related to the accuracy or integrity of any part of the work are appropriately investigated and resolved. All authors qualify for authorship and all those who qualify for authorship are listed.

## Acknowledgements

Image processing and analysis for this work was performed in the Confocal and Specialized Microscopy Shared Resource of the Herbert Irving Comprehensive Cancer Center at Columbia University, supported by NIH grant #P30 CA013696 (National Cancer Institute). Additionally, we are grateful to other members of the Bartoli Brain Tumor and Schevon Lab for their useful comments.

## Data Availability

The data sets are available from the corresponding author on reasonable request.

## Supplementary Information

**Supplementary Figure 1.**
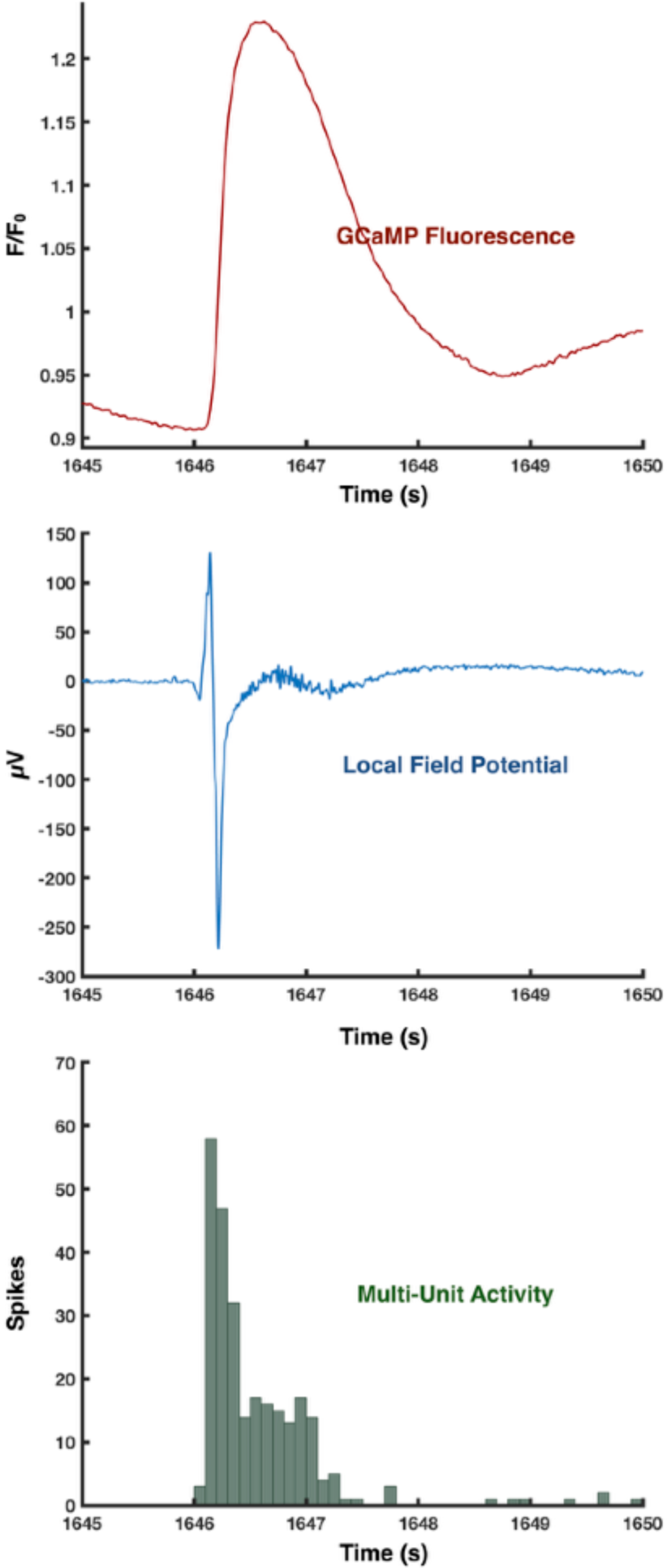
High resolution view of GCaMP fluorescence, LFP, and MUA acquired from the same electrode in a tumor-bearing slice.

## Supplementary Video 1

**Wide-field optical imaging of an untreated tumor-bearing slice during an ictal event in the presence of zero-magnesium solution.** Immunofluorescence micrograph demonstrating the histology for the slice is displayed to the left. Thy1-GCaMP positive neurons (green) and HA positive glioma cells (red) are shown. Cells positive for phosphorylated S6 ribosomal protein are also displayed (magenta). Video is taken at 50 fps.

## Supplementary Video 2

**Wide-field optical imaging of an untreated tumor-bearing slice in response to stimulation at a single microelectrode in aCSF + picrotoxin.** Cyan circle denotes the region of interest (ROI) around the stimulated electrode. Microelectrode stimulation is delivered to electrode 61 at 50 µA. Time of stimulation is annotated “stim” and is also denoted by a change in the color of the ROI to red. Immunofluorescence micrograph demonstrating the histology for the slice is displayed to the left. Thy1-GCaMP positive neurons (green) and HA positive glioma cells (red) are shown. Cells positive for phosphorylated S6 ribosomal protein are also displayed (magenta). Video is taken at 50 fps.

## Supplementary Video 3

**Multimodal recording of an ictal event in a control slice.** Heatmaps of GCaMP fluorescence and LFP during a 40 second segment of a recording from a control slice. Radii of circles (magenta for FS cells contributing to surround inhibition, black for FS cells recruited to excitatory activity) on heatmaps indicate the instantaneous firing rate of the FS cells in that electrode. LFP from a representative channel and raster plot of the FS cell firing shown below.

## Supplementary Video 4

**Multimodal recording of an ictal event in an untreated tumor-bearing slice.** Heatmaps of GCaMP fluorescence and LFP during a 40 second segment of a recording from an untreated tumor-bearing slice. Radii of circles (magenta for FS cells contributing to surround inhibition, black for FS cells recruited to excitatory activity) on heatmaps indicate the instantaneous firing rate of the FS cells in that electrode. LFP from a representative channel and raster plot of the FS cell firing shown below.

## Supplementary Video 5

**Multimodal recording of an ictal event in a treated tumor-bearing slice.** Heatmaps of GCaMP fluorescence and LFP during a 40 second segment of a recording from a treated tumor-bearing slice. Radii of circles (magenta for FS cells contributing to surround inhibition, black for FS cells recruited to excitatory activity) on heatmaps indicate the instantaneous firing rate of the FS cells in that electrode. LFP from a representative channel and raster plot of the FS cell firing shown below.

